# Big Behavioral Data: Psychology, Ethology and the Foundations of Neuroscience

**DOI:** 10.1101/006809

**Authors:** Alex Gomez-Marin, Joseph J. Paton, Adam R. Kampff, Rui M. Costa, Zachary F. Mainen

**Author notes:** To whom correspondence should be addressed at, Champalimaud Neuroscience Programme, Champalimaud Centre for the Unknown, Avenida de Brasilia, s/n 1400-038, Lisbon, PORTUGAL.

## Abstract

Behavior is a unifying organismal process through which genes, neural function, anatomy and environment converge and interrelate. Here we review the current state and sketch the future impact of accelerating advances in technology for behavioral studies, focusing on rodents as an exemplar. We frame our analysis in three dimensions: (1) degree of experimental constraint, (2) dimensionality of data, and (3) level of description. We argue that ethomics, fueled by “big behavioral data”, presents challenges proportionate to its promise and describe how these challenges might be met through opportunities afforded by the two rival conceptual legacies of 20^th^ century behavioral science, ethology and psychology. We conclude that although “ethomes” are not the goal, big behavioral data has the potential to transform and unify these two disciplines and to solidify the foundations of others, including neuroscience, particularly if the data is as open as it is copious and quantitative.

## 1. Introduction: Behavior is foundational

Behavior is “what animals do”. It can be defined as the muscular output of an organism or, alternatively, as the externally observable dynamical features of an organism (Box 1). The brain is the chief architect, orchestrator and driver of behavior. Leaving aside purely subjective states, behavior is conversely the principal function of the brain, as respiration is the function of the lungs. If the problem of neuroscience is to understand brain function, then success hinges not only on explaining how neural systems work but in linking this to behavior in a systematic way. For these reasons, behavioral observations may not always be at the forefront, but are inevitably to be found at the core of brain science. Where respiration is not so taxing to describe, behavior is as complex as the nervous system itself. Yet the collection and systematization of behavioral data is obligatory to give meaning to neural data, and any other “omics” will ultimately miss the very point of the brain without this foundation ().

### Box 1

#### Defining behavior and its key features

Animal behavior is the macroscopic expression of neural activity, implemented by muscular and glandular contractions acting on the body “plant” and resulting in *egocentric* and *allocentric* changes of the animal in an organized temporal sequence. While we focus on rodents, behavior across all species is an expansive concept, ranging from speech, gestures and writing, to micropostural adjustment, reaching and locomotion, from facial expressions, sneezing and crying, to flying, diving and sonar emissions, not to mention construction of burrows, webs, buildings and bibles. Three key attributes of animal behavior are:

1. Behavior is **relational** to the environment. The relationship of the animal to its environment (including other animals) defines affordances (opportunities for behaving). These are needed for understanding and explaining behavior. This implies the need to specify the context—environment or “assay—in which behavior is defined.
2. Behavior is **dynamic**. As physiology is distinguished from anatomy by dynamics, behavior is manifested through time. Thus, frameworks for time series analysis are critical to all behavioral analysis. Even to speak of “a behavior” as opposed to “behavior” implies a chunking of this time series that is itself a non-trivial inference.
3. Behavior is **high-dimensional**, complex and variable (unpredictable). The number of behavioral effectors and their degrees of freedom (e.g. arm or vocal articulation) reduces somewhat the dimensionality compared to the brain itself, but the number is large enough that we cannot even clearly enumerate it. Bodies limit the simultaneous expression of incompatible behaviors (e.g. go left implies not go right), but do not rule out simultaneous expression of multiple “behaviors” (e.g. talking and walking).

Neuroscience refers specifically to the brain and has as its ultimate goal understanding “how the brain works” or, as it is routinely phrased in the opening lines of countless papers, to understand “how the brain produces behavior”. In their pursuit of a tractable problem, neuroscientists have tended to reduce the complexity of behavior by favoring highly-constrained experimental preparations that allow them to focus on the complexity of the brain itself. Reducing complexity is a requisite of scientific progress, but one could claim that neuroscientists might not even know if they have succeeded in understanding “how the brain works” unless they can also characterize the phenomenon the brain is produces, i.e. behavior. Thus, behavioral data is not simply a tool for helping neuroscientists interpret brain data, but also the foundational problem of neuroscience, one that deserves primary importance for at least three reasons:

First, natural selection acts on behavior, not directly on genes and neural firing patterns. Even knowing all possible details about genes or neurons would be incomplete if we could not relate those spaces to behavior.

Second, behavior is a hard problem. Behavioral repertoires are large and complex; they overlap with no clear separation of scales. Further, it is not the brain alone that produces behavior, but rather its interaction with an even more complex environment.

Third, behavior is not merely the output of the brain, it is the unifying space where genes, neural structure, neural function, body plan, physical constraints and environmental effects converge (Figure 1). In fact, behavior is a natural continuum in which some of the most challenging questions of physics, biology and psychology, and the social sciences converge.

**Figure 1.**
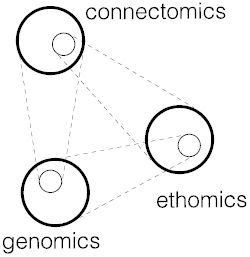
Big data in the context of connectomics and genomics. The putative relationship between behavior and other “spaces” (i.e. genes, neural function, neural connectivity). The illustration is adapted from Penrose’s triple world representation and implies a paradox or at least a loop: behavior emerges from connectivity, connectivity in turn is dictated by the genome (and environment), and gene selection is in turn determined by behavior.

The study of behavior has a long and rich history that we must try to summarize in order to frame our view of the future. Darwin proposed that behavior is selected through evolution^1^, implying that behavioral units or patterns are encoded biologically and expressed in future generations (comparable across individuals) and in closely related species (comparable across species). In the 19^th^ and 20^th^ centuries, two main lines of approach brought us to the modern age.

On one side, the ethologists developed efforts to understand behavior in natural environments^2–4^. Ethologists sought principles of the organization of primarily innate behaviors^3^, common rules governing behavior across species. This led to concepts such as imprinting and releasing mechanisms^2,3^. They also developed methods to define how behavior patterns are composed of simpler parts (ethograms) and sought to describe the whole behavioral repertoire of a species, what we might call “ethomes”.

In a second stream, mainly within physiology and psychology, behavior was studied in less natural and more controlled “laboratory” settings. These schools, including the “behaviorists”, developed paradigms primarily focused on learned behaviors relating stimuli, actions and outcomes, including classical conditioning^5^ and instrumental or operant conditioning^6^. Their search for general principles of learning and motivation led to the development of principles including drive satisfaction^7^, the generation and selection of behaviors based on their consequences^8,9^, the formation and use of cognitive maps by which novel solutions can deduced from experience^10–12^.

We stand on these giant shoulders with a sense of progress, but without a glimpse of the horizon. Where are we? From the similarities and differences between these prior efforts, we can define three primary axes in which to consider the goals, limits, and future opportunities for behavioral studies (Figure 2):

1. The degree of **constraints** imposed by the experimental context—lower in the case of ethologists, high in the case of the psychologists. Constraints limit the scope of behavior, narrowing the possibilities for expression to a particular set of conditions. Constraints also dictate the affordances^13^ available to the animal required for expression of behavior (e.g. climbing could only be expressed in an environment that affords climbing). Constraints also give a “frame of reference”. Thus, constraints express hypotheses about what is important to observe. A behavioral assay, by standardizing constraints, facilitates comparison and replication of behavior within the same assay, but hinders generalization of the findings to new settings.
2. The **level of description**, ranging from the complexity of primary data (low level) to abstract metrics and general concepts (high level). One can analogize this to way neuroscience pictures the brain as moving from primary sensory representations to view invariant object representations. General principles are the goal of the scientific process (and of vision) and this was shared amongst all behavioral schools. Nevertheless, applying high-level descriptions narrow and blind the experimenter to other interpretations. A child drawing a chair reproduces not what hits his retina, but his concept of a chair (four legs, etc.); she is blind to its true form.
3. The **dimensionality of data**, ranging from low to high. Here, classical ethologists and psychologists were both limited to lower dimensional data available by direct observation note-taking, etc, thus limiting the precision and dimensionality of their records. The development of computer and information technology has opened up new possibilities, offering us bigger, more precise, higher dimensional observations.

**Figure 2.**
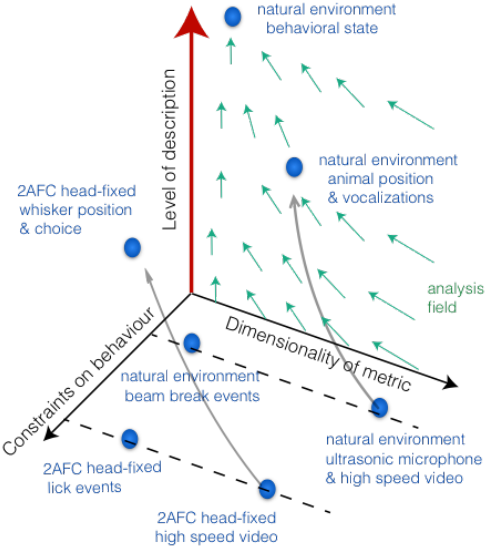
Behavioral data systematization in three dimensions. We conceptualize behavior along three axes: (1) the constraints of the experimental environment, ranging from few (more natural) to many (more controlled); (2) the dimensionality of the measurements, ranging from low to high (as implied by “big data”), and (3, in red) the level of description or explanation, from low (concrete and specific) to high (abstract and general). Blue dots represent several different behavioral paradigms illustrating where they fall in this space. The gray arrow field illustrates the goal of behavioral science: to move from low to high level descriptions (up the third axis) by reducing the dimensionality of the data.

Within this conceptual framework, we can now depict the legacy of previous studies, the promise of “big data” and the challenges faced in applying it (Figure 3). We can see (and will argue) that (1) the issue of constraints is an old conceptual struggle still unresolved, (2) moving to higher levels of description remains a universal scientific motive, but one that is also dangerous when premature, and (3) technology delivering big data has enlarged the “playing field” in ways that interact with (1) and (2).

**Figure 3.**
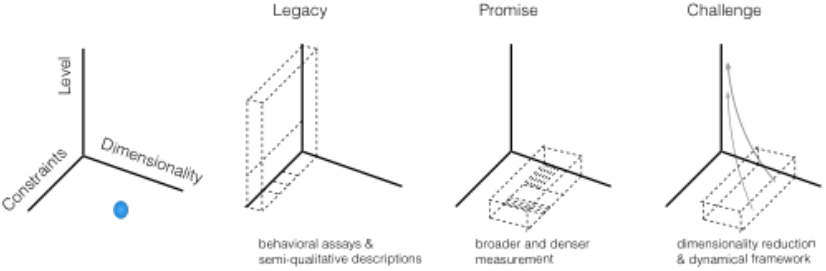
The behavioral space and its relationship to the legacy of psychology and ethology, the promise of big behavioral data, and the challenges. **Legacy**: the achievements and limits of the tradition: several particular assays that link low dimensional measures -sometimes qualitative- to higher levels concepts. **Promise** of big behavioral data: to sample more densely in a systematic fashion and to interconnect different sampled regions by means of standardizing and sharing data. **Challenges**: to use carefully chosen constraints (hypotheses) to move from high-dimensional to low dimensional and from low level to high level descriptions).

The primary aims of this review will be to articulate whether and how “big data technology” has changed the landscape of behavioral studies. We will consider the implication of this for (A) the development of behavioral assays (constraints), particularly the balance between more and less constrained assays and (B) the possible need to revise old behavioral concepts and the possibility of paradigm shifts. We will do so through the primary lens of rodent behavioral studies, whilst bringing comparisons to other species and other “omics” when useful. Throughout, we will stress the implications of these behavioral issues for the interpretation of neural data. We first (Section 2) review the tools and technology that are contributing to “big data” and the opportunities for behavioral studies that they create. We then (Section 3) discuss the challenges that exist in applying those tools in a scientifically productive manner. In the following two sections (Sections 4 & 5), we offer more specific vision of how big data approaches can transform modern versions of the “psychological” and “ethological” approaches to rodent behavioral analysis. We conclude (Section 6) with a sketch of the longer-term promise of the “big behavior” for the future of neuroscience.

## 2. Promise: Advancements in technology and a new era of “big behavioral data”

The study of behavior has long benefitted from advances in technology that have improved both the quality and quantity of the data that can be acquired. Currently, tools developed for other applications (e.g. video games, media streaming, computer vision, etc.) have made it possible to inexpensively acquire, store, and analyze vast amounts of behavior data, often in an automated manner. In contrast with other “omics” projects (e.g. genomics or connectomics), the acquisition of a large “ethomics” dataset is technically straightforward; what data these datasets should contain, such that they will advance our understanding of how brains function, is still unclear and will be the main topic of this review. However, we begin by highlighting some of the new possibilities enabled by current technology and how they enable access to new domains of behavioral investigation.

Human observation has been the standard approach to studying behavior for millennia. The advent of technology for acquiring records of these observations (photography, videography, etc.) found immediate application. These new approaches provided not just a tool for documenting observations, but also offered access to new spatial and temporal scales that were inaccessible to an unaided human observer^14^ (e.g. high-speed video, ultrasonic microphony, infrared illumination, etc.). Such “augmented observations” have become vital to many domains of behavioral neuroscience, yet they have historically been the pursuit of specialists requiring sophisticated, expensive equipment. Fortunately, driven by consumer interest in documenting the behavior of their children, cats, and extreme sports exploits/mishaps, this technology has become much more accessible. Such advances have made it feasible to acquire, cheaply, extremely detailed audio and video records of animal behavior, continuously, for an entire experiment. Currently a full video record of a rodent’s lifespan (∼24 months) in a standard cage (mm resolution), at VGA resolution, day and night (with an IR LED illumination) and lossy yet sufficient compression, can be acquired with a $50 webcam and stored on a hard drive.

Such “augmented” behavioral observations are not restricted to standard media (audio, visual) modalities. Sensors developed for smartphones, e.g. accelerometers, gyroscopes, GPS, etc., can provide new measures of behavior in a robust, miniaturized package. These sensors, which were designed to operate at low power in wireless devices, can be affixed to an animal, and transmit detailed, continuous measures of behavior over long times periods. Inertial sensors (accelerometers) have been used to extract continuous acceleration data from bodies of animals, including humans, with high temporal resolution and long durations^15–18^. Furthermore, measurements from different types of sensors can be combined to infer more accurate measures^19^. For example, high temporal resolution velocity and position data can be computed by integrating acceleration, if these egocentric measures are then be combined with a geocentric reference, provided by a video or GPS data, and corrected for sensor drift, then, in theory, inertial sensors could replace the use of video for animal tracking, but working systems are not yet commercially available. The same microprocessors that make wireless transmission of inertial data possible also permit the sampling of other data sources. Any signal that can be turned into an analog voltage can be mated with a wireless system. Thus, continuous wireless sensing of behaviorally-relevant physiological signals such as body temperature, respiration via pressure or nasal temperature^20^, heart rate, and electromyogram (EMG), as well as with neural recordings^18,21^.

Historically, a major constraint in the amount of behavioral data acquired has been the human resources required to perform each experiment. Advances in technology have made it increasingly feasible to automate the behavioral assay and context and thereby collect more data from a given animal and/or more animals, important issues to which we will return (and see Box 2). Automation of assays affords the substantial advantage of greater inter-lab reproducibility and the possible disadvantage of commitment to a given set of constraints implied by a uniformized behavioral setup. These approaches have already seen substantial application in smaller species, but have already proven valuable for rodent studies as well (reviewed in ^22^). Two levels of automation have been deployed to ‘scale up’ behavior. First, systems in which smart software reduces the need for human monitoring and intervention or allows it to be done remotely or offline^23^. These systems still require manual transfer of animals into and out of cages, etc. Second, and more radically, are “live-in” systems in which rodents live in the behavioral assay or shuttle between “home cage” and assay by themselves^24^. Commercial systems have integrated RFID technology with sensors and actuators (e.g. food and water dispensers) to create complex environments where multiple types of data can be collected over days, demonstrating dependency of behavioral phenotypes on geotypes and genetic manipulations. Other commercial systems have used high-resolution video and automatic classification to characterize the order and frequency of behaviors, revealing pre-symptomatic behavioral deviations in mouse models of disease^25–27^. Yet the more recent development of far less expensive hardware (e.g. Arduino microcontrollers) and open software is a potential game-changer.

### Box 2

#### Big open data—options and imperatives

The “problem” of behavior is difficult, and the more people involved in tackling the problem, the better. It is now possible to share raw behavioral data, yet we lack accepted standards to efficiently and productively do so. Ought experimenters be compelled to record store and share their raw behavioral data, just as they are being compelled to share, e.g., genomic data?

**Con**

- Collecting data is not equivalent to doing experiments. Experiments require well-conceived designs that probe particular aspects of behavior; simply generating large datasets, and sharing them, risks diluting such efforts.
- Storage and sharing is always becoming cheaper, but it is still not negligible.
- Each lab has their own particular assays and conditions, making comparisons difficult even if the data is openly available. The resulting confusion could impede collective progress.

**Pro**

- Storing and sharing primary behavioral data would allow researchers to revisit it, even much later, in search of, or in light of, new insights.
- Along with automated behavioral assays, open data and shared analysis tools will facilitate comparison of data across laboratories.

**Suggestions**

- Data sharing standards will only arise in an environment that requires them, and it should be acceptable to request primary behavioral data from the source lab.
- It need not, yet, be required to make all raw data available, as the challenges of such a mandate might overwhelm the more pressing need for innovation.
- Researchers who manage to both acquire and share such data should not only be encouraged but rewarded.
- We ought to encourage different solutions to these problems, identify what works and facilitate its adaptation.

Although our main focus in this review is on the laboratory-based behavior of individual animals, behavior evolved in the natural environment and experiments in the laboratory have sampled a greatly reduced subset of behaviorally relevant environments. Efforts to export quantitative methods to more ecological situations are also taking advantage of advances in technology, opening up many new possibilities for monitoring freely behaving animals in wild or semi-wild conditions over large spatial and temporal scales. Fruitful directions that have being explored include image-based tracking^28^, animal-attached remote sensing, e.g. using RFIDs^21^, autonomous recording tags, animal mounted video cameras, and specifically for terrestrial animals, biotelemetry of physiology as well as location and fingerprints for phenotype recognition and profiling of behavior of individuals and species. Furthermore, the same methods are often also applicable to social interactions that involve more than one animal.

Essentially, regardless of setting, big behavioral data implies that ability to collect and manage large volumes of data both in density and in extension within the behavioral space (Figure 2). By density we mean to move along the “dimensionality” axis, collecting information at higher rates, higher resolution and from more modalities. However, higher density also implies performing measurements for very long time periods, beyond a few minutes and up to hours, days and even the whole life of an individual; and to bring the number of animals tested to a level where one could start asking questions about individual variability (akin to personalized genomes vs. “the” human genome). By “extension” we mean the ability to explore more thoroughly the “constraints”. High capacity for measurement means one could sample many different behavioral assays and, if the data is shared, to be able to compare across tasks and laboratories. Therefore, in a weak sense, big data means more precision, more resolution, longer observations, higher number of animals, and datasets across a larger variety of tasks and conditions. In a strong sense, big data suggests the possibility of “fully mapping” the behavioral space, and thus it implies the possibility of the so-called “ethome” (in analogy with the genome or connectome) (Figure 3). We return to this issue in detail in Section 5.

Similar to other fields dealing with massive data sets, reduction of dimensionality in behavioral calls for algorithmic solutions. Fortunately, methods in statistics and machine learning developed for solving similar problems in various domains (e.g. internet search, artificial intelligence, machine vision, linguistics, etc.) have enormous applicability to behavioral data (see review, this issue Nature Neuroscience). We do note that, however, with great power comes great responsibility—to understand what these solutions are doing and not treat them as black boxes.

While acquiring high dimensional data and then compressing it again, it might seem that we are back to where we started. But while a manual observer scoring is irreversible, a video or sensor recording can be “reinterpreted” using different conceptual structures. Imagine we had access to videos of Tolman and Lorenz’s original experiments. Thus, the possibility to record systematically higher dimensional data might provide new insights by escaping the assumptions of previous experiments. In this process, aspects of the data that were previously marginalized or deemed as noise might reveal themselves as important biological dimensions that further clarify and amplify our explanatory power. This potential would be maximized by efforts to encourage data sharing, although this issue is not without complexities (Box 2).

To fulfill the promise, how exactly are we to use these advantages to the maximum? What are the challenges of large amounts of higher-dimensional behavioral data, from cheap, automatized experiments for the design of behavioral assays and for the possible reconceptualization of behavior itself?

## 3. The challenges of big behavioral data

It is clear that we have the capacity to acquire big behavioral data, but the main challenges lie ahead. A ballpark estimate of the dimensionality of the “raw” data from a “manual” ethogram would be <100 behaviors at a time resolution of <1 Hz. Assuming that each “behavior” is itself an independent one bit channel of information, that yields a max data rate of 100 bit/s, or about the data rate of a single inertial sensor axis. Contrast this to a video at 1 megapixel, 8 bit frames at 120 Hz that represents a bandwidth of 1 gigabit/s. This yields an astonishing 10-million (10^7^) fold increase in data rate. Even allowing for 1000-fold compression to a “mere” 1 megabit/s, something is clearly unfair about this comparison. The “manual” pre-video scoring has taken the “raw data” on the observer’s retina and converted it into much higher-level abstract concepts (e.g. “the rat froze”). What is being recorded is not merely lower-dimensional, it is also “higher level” (Figure 2). The video camera captures data, but it is not meaningful until it is “processed” and its dimensionality reduced. A complete description or library of the behavior of a particular individual or species under certain circumstances is a complex but perhaps not impossible goal. But a raw video library, however exhaustive, is a long way from this.

Prediction and understanding require more than data collection, they require synthesis and reduction of data to “principles”. In the behavior space we have outlined, we need to reduce the dimensionality of the data whilst moving from lower level to higher level descriptions (Fig. 1). To do so requires using, creatively and insightfully, constraints that are implied by the choice of behavioral context or assays and analysis techniques. In other big data projects, the “units” that needed be measured and the conceptual framework to structure and analyze the data were established a priori. For genomic data, it was known that one ultimately needed to read strings of nucleotides (unless our framework is epigenetics); for electrophysiological data, we know we need to extract spike times (unless our framework gives emphasis to local field potentials); for a connectome, all synaptic partners must be identified (unless our framework relies on knowledge of synaptic strength). For behavioral data, there are many additional challenges that will call for the application of conceptual frameworks and many issues that are not solved by high capacity data recording and analysis alone. Consider among these:

### Difficulty of time segmentation

Because behavior unfolds in time, “segmenting” or “parsing” it into discrete chunks is a common (though not universal) step in both psychological and ethological approaches. But segmenting behavior will be conceptually challenging if temporal organization is hierarchical or multi-scale. For example, one can differentiate and classify locomotion episodes and grooming episodes, but since both are rhythmic behaviors, one could further segment each step or grooming cycle as smaller repeated elements.

### Poverty of environmental stimuli and affordances

As we have discussed, behavior happens in, and because of, the environment. Big data recording and analysis may bring behaviors into view, as a telescope makes visible a universe not visible to the naked eye. But unlike for the astronomer peering into the night sky, without providing equally rich stimuli and affordances in the environment, even the most detailed video recording and analysis can only capture a tiny fraction of an animal’s “ethome”. This point is illustrated by sensory neuroscience studies, which by exploring “stimulus space” probes the dependence of an animal’s simple binary responses on the environment. How do we even know what the relevant stimulus space is? While some kinds of stimuli can be computer generated, some cannot (e.g. odors). The problem is even more acute for the side of action itself, which, lies in a physical world that is only sparsely computer controllable. For example, nothing yet will substitute for an encounter with a conspecific or predator. How can “big data” approaches deal with these issues?

### Limits of controlability

It is also important to consider that the environment constrains but does not strictly limit the expression of behavior. That is, animals are still left to a larger or smaller degree with variance in many axes. Compared to our ability to record what is being emitted, our ability to strictly control those variables remains vastly limited. This problem can be compared to that faced in multi-neuron data, where despite advances in optogenetics, our ability to record (in terms of numbers and temporal precision) still vastly outstrips our ability to control. One could contrast this with molecular biology, in which progress depended not on the ability to measure but on the ability to write (cut, paste, construct) genetic information in the relevant space.

### The conundrum of standardization

Although standardization, promoting automation, is fundamental to the collection of “big data”, it presents an awkward dilemma. The brain evolved to control behavior in a complex, rich, and uncontrolled environment, and many of its most remarkable functions are those related to its ability to adapt to these diverse demands and changing conditions. If our efforts to produce “big data” result (or require!) avoiding complexity in the experimental environments of our assays, then it is unclear if this data will ever be able to inform our understanding of the brain’s most impressive capacities.

As we have argued, these challenges are not without solution, but to solve them we will need to rely on conceptual frameworks that determine what features of behavior are meaningful. Most frameworks don’t yet make any sense of the myriad features that might be discerned in video. For example, classical learning theory from psychology do not make strong predictions about say the details of the articulation of the arm when pressing a lever. Other frameworks, such as optimal control theory combined with biomechanics do make predictions about the articulation of the arm, required joint torques, etc., yet can predict little about arm behavior in a food foraging task.

The lack of a consensus framework means there is no universal solution for “annotating” behavior or reducing its dimensionality. Each such method implies choices. As we have seen from the legacy of psychological and ethological approaches, there is a tension between relatively constrained and unconstrained approaches. Therefore, we will examine the impact of “big data” on two different approaches, first (Section 4) “decision-making and reinforcement learning”, which is closely aligned and inspired by classical psychological approaches and second (Section 5) “modern ethology and state space analysis”, more closely aligned with ethology approaches to innate behaviors.

## 4. Big behavioral data in the framework of decision-making

To see how decision-making studies can make use of “big data” technology whilst meeting the challenges presented, we consider first the “legacy” of these approaches (Figure 3). We use the term “decision-making” to refer to two closely related approaches. The first is perceptual decision-making or psychophysics, a classical approach to quantitatively linking physical characteristics of stimuli with their perceptual impact^29^ that has been used in rodents to characterize perceptual and cognitive processes^30,31^. The second is reinforcement learning (RL), an approach mainly concerned with learning how to act in a given situation in order to maximize reward or value^32^. As perceptual thresholds and sensitivities are measured in psychophysics, state value functions are inferred from patterns of choices, with more frequent choices reflecting states or actions of greater value^33–35^. Both build on classical psychological work on reinforcer-driven learning (i.e. rewards and punishments) to guide behavior allowing the experimenter to isolate, exaggerate, and systematically explore behavioral functions. This methodology facilitates a mechanistic understanding in which the parts of systems are isolated and manipulated.

### Opportunity 1: Scaling up

A major benefit of “big data” arises naturally from scaling up of the size of the data set. Decision-making tasks in rodents are much more powerful when they include the ability to collect many 100’s to 1000’s of trials in a single session or to amass 10’s or even 100’s of thousands of trials from a given animal or a behavioral data set. The ability to apply relatively “high-throughput” automated (e.g. live-in cage)^24^ or semi-automated behavioral assays can greatly aid in reaching these levels of trials. Such a large corpus of data may reveal aspects of behavior that are small but lawful and those that may require conditioning on many different variables (i.e. dividing the data set amongst many conditions). This is particularly important in analyzing the effects of learning in past trials on the performance of a given trial^36^, because the number of conditions to be included increases exponentially with each past trial. As a simple example, if there are 8 stimuli to be considered and 2 possible choices, there are 16 current trial types, but 256 trials types considering all combinations of current trial and preceding trial. Having very large data sets that allow conditioning on this history may reveal stimulus/choice history effects that would otherwise have been simply choice variability.

### Opportunity 2: From discrete to continuous measures

The core unit of many psychological assays is the “trial”, which solves the “temporal segmentation problem”. Trials are composed of several different phases in a sequential chain. For example, a trial might begin with a criterion of the animal signaling its readiness and then proceed from stimulus presentation to response to outcome. An experimental session typically includes rules about how longer sequences of trials are structured (e.g. different types of stimuli randomized or clustered in blocks, possibly changes based on performance criteria, etc.).

We believe big data approaches can benefit decision-making studies by expanding the dimensionality of the measurements being made. While thus far most of these approaches have relied mainly on minimal binary response measures (e.g. lever up, lever down; infrared beam breaks with the snout), the magnitude of internal perceptual or cognitive variables are likely to be continuously valued and evolving in time. Because in neural terms “motor systems” are not fully insulated or isolated from the “cognitive systems”, information about the time course of unfolding decisions may be found in continuously expressed behavior such as the micromovements of the head^37–40^.

### Opportunity 3: Noticing the unconstrained

In decision-making approaches it is traditional to constrain the available behavioral outputs as tightly as possible. Rodents may indicate choice through selection of a particular physical path while moving through a maze^37^, one of a number of available nose ports^20,23,30,41^ or levers, or by positioning a manipulandum or licking at particular reward delivery tubes. One of the biggest impacts of “big data” approaches will be to reveal the “unconstrained” as a rich source of insight rather than a nuisance. Consider that decision-making studies principally reinforce and measure binary choice output (e.g. left vs. right), but it has long been known that response times, when unconstrained, are extremely revealing about the behavior^42^. More recently, is has been seen that free response time reveals much more as well. For example, response times to obtain outcomes reveal expected value^43^, allowing one to measure value on a trial-by-trial rather than average basis, and the waiting time of an animal for a delayed reward indexes confidence in the preceding perceptual decision^31^. In a similar manner, the application of high-speed video data is likely to add key additional insights for the dynamics of less constrained movements executed in the course of meeting task demands^41^.

### Opportunity 4: Virtual and augmented reality

Computers have already made exploring stimulus space a relatively tractable problem, at least in the visual and auditory domains, while somatosensory, vestibular and olfactory and other domains remain much more challenging. When considering the experimental subject not only as being acted on, but acting on the world, the issue is also challenging for computers. But here we believe there is also a very interesting opportunity for big data in the psychological framework. Once acquiring a rich measure of behavior (i.e. video) it will also be powerful to define, “virtually”, quantitative readouts. For example, in the classic operant conditioning lever-press paradigm, a more flexible reporter could be implemented using a video-based real-time feedback control system. Approaches like this have already been implemented with respect to location in locomotor behavior^44,45^, which is undoubtedly an ethologically important domain for rodents. To provide richer opportunities, even more complex readouts can be harnessed to give feedback to the animal, providing “virtual affordances”, e.g. rearing events could be detected and linked to a reward or another stimulus. This approach could also provides a natural connection to more ethological descriptions of behavior.

### Opportunity 5: Computational models and big data

A key feature of successful studies of more complex phenomena by psychological approaches is the use of computational models that provide a mathematical description of the features of behavior. These models comprise “higher level” descriptions of behavior by which raw behavioral data is transformed from a “lower level” and higher dimensional representation by inferring the dynamics of a smaller number of state variables. Three examples are “integrated evidence” in models of bounded accumulation of evidence^39^, “experienced value” in models of reward-based decisions^46^ and “subjective confidence” in models of higher order decision-making^47^. Critically, these models serve as the scaffold by which the results obtained from a specifically constrained behavioral assay can be generalized to other assays and environments. More detailed models instantiate abstract concepts (theories) in a way that allows them to be cached out into observables. Critically, in addition to providing concise and predictive models of behavior, these models also constitute “linking hypotheses” through which behavioral and neural data can be related^48,49^.

Big data may have its biggest impact when applied to computational models that capture (attempt to predict) behavioral features. This is because the severe problem of existing computational models is that they easily become too complex and therefore underconstrained. The more complex a model is, the larger and richer a set of data that is needed to test or constrain it. The marriage between “big behavioral data” and complex brain/behavioral models, which is still not yet in its honeymoon phase, is likely to be a long and rich one.

## 5. Big behavioral data in the framework of ethology

Ethology, the study of animal behavior under natural conditions, relied on extensive observation and annotation of behavioral states and events by human observers. Particular attention was often paid to the phylogentic history of the species being studied and the selective pressures applied to organisms by their natural and social environments. The legacy of modern neuroethology, specifically when compared to the psychological approach, provides less constrained experiments: rather than placing constraints in the environment and creating special places by building levers and pokes, ethologists let animals express their behavior more freely at the expense of control (Figure 2).

### Opportunity 1: Making ethograms reversible

The annotations of expert ethologists grasped meaningful behavioral states by segmenting the continuous flow of behavior into a sequence of discrete categories sewn together by transitions between those states in an ethogram representation. Segmentation embodies the theory in the observation^50^, which is irreversible: an ethogram cannot be reversed into the underlying phenomena (Figure 1; a floor on the levels axis that would not allow going back to lower level descriptions. Starting with and conserving high-dimensional measurements allows to different routes to moving up the z-axis, as illustrated by the green arrow operators). Computer-based approaches transform the ethological approach because data acquisition is no longer inextricable from data analysis. Now, with big behavioral data, continuous high-resolution multidimensional streams of raw data can be collected, saved and shared, thus providing the opportunity to revisit the raw data as many times as necessary, without getting stuck in ad hoc (pre-defined behavioral atoms) summary statistics. Data can be shared and reanalyzed by the same or different laboratories. As annotation remains laborious even with computers, data can also be collected and stored “just in case”, allowing inspection in later stages of an experiment.

### Opportunity 2: Scaling up in effort and timescales

Ethological approaches rely on behavioral classification or annotation to identify features of interest and quantify their occurrences and relationships. Big data approaches provide the opportunity to automate this by the use of computer algorithms. This process can be supervised, that is aided by the judgments of trained observers, in which case this judgment of the scientist is translated into an algorithm that then prescribes the processing of the data^51^. It can also be unsupervised, relying on features of the data itself for classification, and potentially less biased^52–54^. Automated annotation allows fixed rules to govern segmentation of behavior over large amounts of data, providing standardization. It is also vastly faster, therefore greatly expanding the scale of what can be annotated. This opens the opportunity to study the behavior of individuals at very short and very long time scales previously inaccessible to unaided observers^55,56^. From ultra-fast maneuvers during prey capture^57^ to non-rapid behavioral assays in naturalistic conditions^58^, big data approaches can provides a window into entirely new phenomena.

### Opportunity 3: Finding simplicity in higher-dimensions

Big data makes it possible to densely sample the many degrees of freedom of a behavioral process. While segmentation and ethograms are one way to look for simplicity in higher-dimensions, by avoiding premature coarse-graining or segmenting of data it is possible to apply many other fundamental theoretical frameworks that operate on high dimensional continuous data itself, such as information theory^59,60^. For instance, applying a statistical mechanics formalism, the collective behavior of flocks of birds, measured through continuous correlations in the location and velocity across neighbors, is seen to be posed at criticality^61^. Beyond the ethogram representation, big data allows one to densely estimate the distance between distributions of continuous kinematic variables^62^, the covariance matrix of body postures^63^, motifs from times series and grammars^64,65,66^, or the low dimensional embeddings naturally emerging from spatiotemporal patterns in pixel space^67^, thus mapping the phenotypic space^68^.

### Opportunity 4: Contrasting contexts: sampling, quantifying and recreating the Umwelt

A fundamental mission of the ethological approach is to integrate the study of the animal behavior with its world or “Umwelt”^69^. Thus, thoroughly characterizing an animal’s natural behavior generally requires observation across different environmental conditions. Big data approaches will facilitate this both by enabling larger-scale studies and by opening new possibilities such as monitoring and control the sensory input as an animal negotiates its world (e.g., estimating the temporal dynamics of olfactory input as it orients in chemical gradients^70^). Exploring behaviors across natural environments from the animal’s perspective is very revealing^71^. One can examine the relationship between sets of different behaviors to determine which are invariantly associated vs. those that are merely coincidental^72,73^. Similarly, cross-environment comparisons will help to distinguish circumstantial from essential neural-behavioral correlations^74,75^. Finally, contrasting apparently similar behaviors in different environments can also reveal different causes of apparently identical behaviors. For example, even careful examination of the movement dynamics of lever pressing in rodents is unable to reveal the difference between habitual and goal-directed that can be distinguished by environmental manipulations such as sensory specific satiation and contingency degradation. Thus, environmental manipulations can be critical to understand “why” a certain behavior is being performed, revealing alternative neural substrates for the “same” action^76^.

### Opportunity 5: De-aggregating variability

Big data approaches will greatly increase not only the size and richness of datasets from each individual, but will bring the number of animals tested in the same assays to hundreds or even thousands, as has been achieved with insects^77^. This combination will allow behavioral descriptions that go from average species behavior to individual behavior. This will allow neuroscience to address the important question of “personality” and individual differences from a neuroethological perspective^78,79^. At the same time, combined with the willingness to share data (big open data, Box 2), this combination of rich data from many individuals will greatly enhance the possibility for identifying sources of variability across laboratories, therefore setting higher standards for experimental protocol/assay design and behavioral analyses and achieving less fragmentary and idiosynchratic descriptions^80^.

### Opportunity 6: Characterizing spontaneous behavioral processes

The stimulus-response approach to behavior has proven as successful as convenient, since one can systematically repeat the same external sensory protocol in order to estimate the statistics of animal responses. In the complementary view, where brains are output-input devices^81^, it is much harder to collect the necessary amount of data to discover high level rules generating the (apparently) noisy behavior^82–85,86^. Big behavioral data will be a key step to expand ethological investigations to the study of spontaneous behavior, where the animal rather than the experimenter calls the shots. Conceiving behavior as a process rather than as a juxtaposition of atoms, one could study behavior using descriptions applied to fluids (e.g. “flow”) rather than solids (e.g. “blocks”). In a continuous dynamical system, transitions “opportunities” emerge from symmetries in the system^87^. From actions never performed before^88^ to the origins of creativity^89^, big behavior will allow us to generate the data necessary for understanding the evolution of behavior^90^, a long standing goal of ethology^91^.

## 6. Conclusions: Ideas of what may come

As we have discussed, the main challenge confronting behavioral science is extracting meaning from ever increasing information. Big behavioral data, despite offering new opportunities, cannot substitute for the development of novel experimental designs and improved conceptual frameworks. When faced with such promise and challenge, behavioral science must define its metrics for achievement. Thus, in closing, we consider what success might look like for behavioral science in the era of big data.

### Richer neural correlates

Recent innovations in imaging and electrophysiology have enabled the collection of increasingly rich descriptions of neural data. However, we risk throwing away as “unexplained” much of that richness for lack of similarly rich behavioral data. Big behavioral data, and the toolset of statistical methods necessary to relate that behavioral data to itself, will provide new approaches to explaining ever greater amounts of neural variability.

### Convergence of ethological and psychological approaches

We can envision a behavioral science based on big-data fostering a unification of the ethological and psychological approaches to animal behavior: the convergence of “trials” and “events”; “biases” and “traits”. For example, three years of continuous monitoring of mice burrowing in the wild allows an experimenter to select from thousand instances of forepaw movement and ask questions that previously were circumscribed to realm of Skinner boxes, all while retaining the essential ethological grounding. Large amounts of data may implicitly contain the conditions necessary to isolate particular behavioral events while controlling for potential confounds, and this is indeed the goal of tightly controlled behavioral paradigms used in the laboratory.

### Ethological homology

Ultimately we seek behavioral universals, elements of convergence that signal life’s phylogenetic and ontogenetic solutions to problems in the physical world^92^. We have thus far a poor grasp of what behavioral universals might look like. For example, an observation that “the rat is rearing” creates a discourse that may be full of assumptions by already assuming universality at the observation stage. Very few studies have succeeded in identifying forms of behavioural invariance and universality across species^93,94^, When considering anatomy, we use the notion of homology across taxa to substantiate the universality of particular forms. A future behavioral science, using analyses of data across taxa, might complete Lorenz’s vision^91^ of establishing homology in ethology.

### Ethomes, sub-ethomes and ethons

One might imagine the product of ethomics be an ethome, a complete description of the set of behaviors manifested by a species in its natural environment. Considering that behavior in a full sense includes complexities such as language and tool use, it is clear that a complete description is impossible for the human species. Even for rodents, as we have argued throughout this review, behaviors must be considered relative to an environment and from a conceptual viewpoint, arguing against a totalistic description. Certainly the pursuit of increasingly detailed descriptions of behavior is useful, and big data approaches are invaluable for exploring a more expansive and previously inaccessible regions of behavioral space. “Sub-ethomes”, descriptions of behavior restricted in a particular environment, may be possible and informative, but a complete description is not the goal. Rather, we would argue that the biggest achievements of “big behavioral science” will be to promote development of new unifying frameworks for addressing behavior. This might lead in the identification of “ethons”, fundamental units of behavior. Ultimately, the success of big behavioral data will foster be the end of its utility: we will then know what degrees of freedom to look for, how, where, and why.

Finally, we want to emphasize that reaching these big goals places an implicit but inexorable constraint on the way we do science: it will demands considerable effort to explore and adopt better ways of standardizing data so that it can be reused and compared. It demands a collaborative relationship with our peers, with an open attitude to share our data and to appreciate theirs.

